# In Breast Cancer ZMIZ1 Co-Regulates E2F2 as Part of the Estrogen Receptor-Mediated Cell-Cycle Response

**DOI:** 10.1101/789610

**Authors:** Weiye Zhao, Susanna F Rose, Ryan Blake, Anže Godicelj, Amy E Cullen, Jack Stenning, Lucy Beevors, Marcel Gehrung, Sanjeev Kumar, Kamal Kishore, Ashley Sawle, Matthew Eldridge, Federico M Giorgi, Katherine S Bridge, Florian Markowetz, Andrew N Holding

**Affiliations:** Department of Biology, University of York, York, YO10 5DD, UK; York Biomedical Research Institute, University of York, York, YO10 5DD, UK; The Alan Turing Institute, 96 Euston Road, Kings Cross, London, NW1 2DB, UK; CRUK Cambridge Institute, University of Cambridge, Cambridge, CB2 0RE, UK; Department of Pharmacy and Biotechnology, University of Bologna, Bologna, 40126, Italy

**Keywords:** Estrogen Receptor, ZMIZ1, E2F2, Breast Cancer, Cancer, Co-factors, Transcription, Nuclear Receptors, Signalling, Prostate Cancer, Patient Outcome

## Abstract

The Estrogen Receptor (ER) drives 75% of breast cancers. On activation, the ER recruits co-factors to form a transcriptionally active complex. These co-factors can modulate tumour growth, and understanding their roles can help to identify new therapeutic targets.

Here, we present the discovery of an ER-ZMIZ1 interaction by quantitative proteomics, and validated by proximity ligation assay. We characterise ZMIZ1 function by demonstrating that targeting ZMIZ1 results in the reduction of ER transcriptional activity at estrogen response elements and a significant decrease in the proliferation of ER-positive cancer cell lines.

To establish a role for the ER-ZMIZ1 interaction, we measured the transcriptional changes in the estrogen response post-ZMIZ1 knockdown using an RNA-seq time-course over 24 hours. GSEA analysis of the ZMIZ1-knockdown data identified a specific delay in the response of estradiol-induced cell-cycle genes.

Integration of ENCODE data with our RNA-seq results identified ER and ZMIZ1 binding at the promoter of E2F2. We therefore propose that ER and ZMIZ1 co-regulate an important subset of cell cycle genes via a novel ER-ZMIZ1-E2F2 signalling axis.

Finally, we show that high ZMIZ1 expression is predictive of worse patient outcome, ER and ZMIZ1 are co-expressed in breast cancer patients in TCGA, METABRIC, and the proteins are co-localised within the nuclei of tumours cell in patient biopsies.

In conclusion, we establish that ZMIZ1 is a regulator of the estrogenic cell cycle response and provide evidence of the biological importance of the ER-ZMIZ1 interaction ER+ patient tumours, supporting potential clinical relevance.

## 1. Introduction

Approximately 75% of breast cancers are classified as Estrogen Receptor (ER) positive. In these cancers, the ER is no longer correctly regulated, subverts cell division regulation, and becomes the driving transcription factor in the tumour (Mohammed et al., 2013). Only a few primary breast cancers have mutations in the ER (Oesterreich & Davidson, 2013), yet the transcriptional activity is frequently abnormal (Carroll, 2016). In many metastatic tumours, the ER is still active and drives the growth of the tumour with a reduced dependence on estrogen or in a ligand-independent manner (Robinson et al., 2013; Toy et al., 2013). This critical role for the ER in disease progression has therefore made the protein a key target for therapeutics like Fulvestrant (Osborne et al., 2004) and Tamoxifen (Jordan, 2003). More recently, Tamoxifen has been prescribed as a preventive treatment in high-risk healthy patients to successfully reduce their chances of developing breast cancer (Cuzick et al., 2015). Most women benefit from endocrine therapy with 50-70% of cases responding to treatment. However, relapse is very common with the risk ranging from 10 to 41% (Pan et al., 2017).

*Co-factors of the ER*. One strategy to overcome relapse focuses on co-factors of the ER. The majority of ER binding sites are at distal enhancer elements (Carroll et al., 2005). On binding to these sites, the receptor catalyses the formation of chromatin loops and recruits several co-activators along with the mediator complex and through these interactions that the ER is able to facilitate the activation of RNA Pol II at the promoters of target genes (Murakami et al., 2017). Without the coordination of the ER and these co-factors it is not possible for the efficient transcription of target genes to occur.

Further characterisation of ER co-factors is therefore essential to identify targets for novel treatment strategies. Successful examples of co-factor based approaches include studies demonstrating that the GREB1-ER-EZH2 transcriptional axis is directly involved in Tamoxifen resistance (Wu et al., 2018) or identifying the pioneer-factor FOXA1 as a key opportunity for future interventions (Nakshatri & Badve, 2007; Behan et al., 2019). This study aims to lay the ground work for future co-factor based therapies by presenting proteomic and genomic evidence that ZMIZ1 is co-expressed with ER in patient tumours and that the inhibition in ZMIZ1 reduces the ability of the ER to promote the cell-cycle within ER+ breast cancer tumour cells via a novel ER-ZMIZ1-E2F2 signalling axis.

## 2. Results

### qPLEX-RIME of ER activation identifies novel co-factor ZMIZ1

We used qPLEX-RIME, a state-of-the-art method for quantifying protein-protein interactions within nuclear receptor complexes (Papachristou et al., 2018), to quantitatively monitor the changes in protein-protein interactions in the ER complex between 0 and 45 minutes in response to activating MCF7 cells (ER positive cell line) with estradiol (Figure 1A). The assay was repeated across four isogenic replicates.

**Figure 1:**
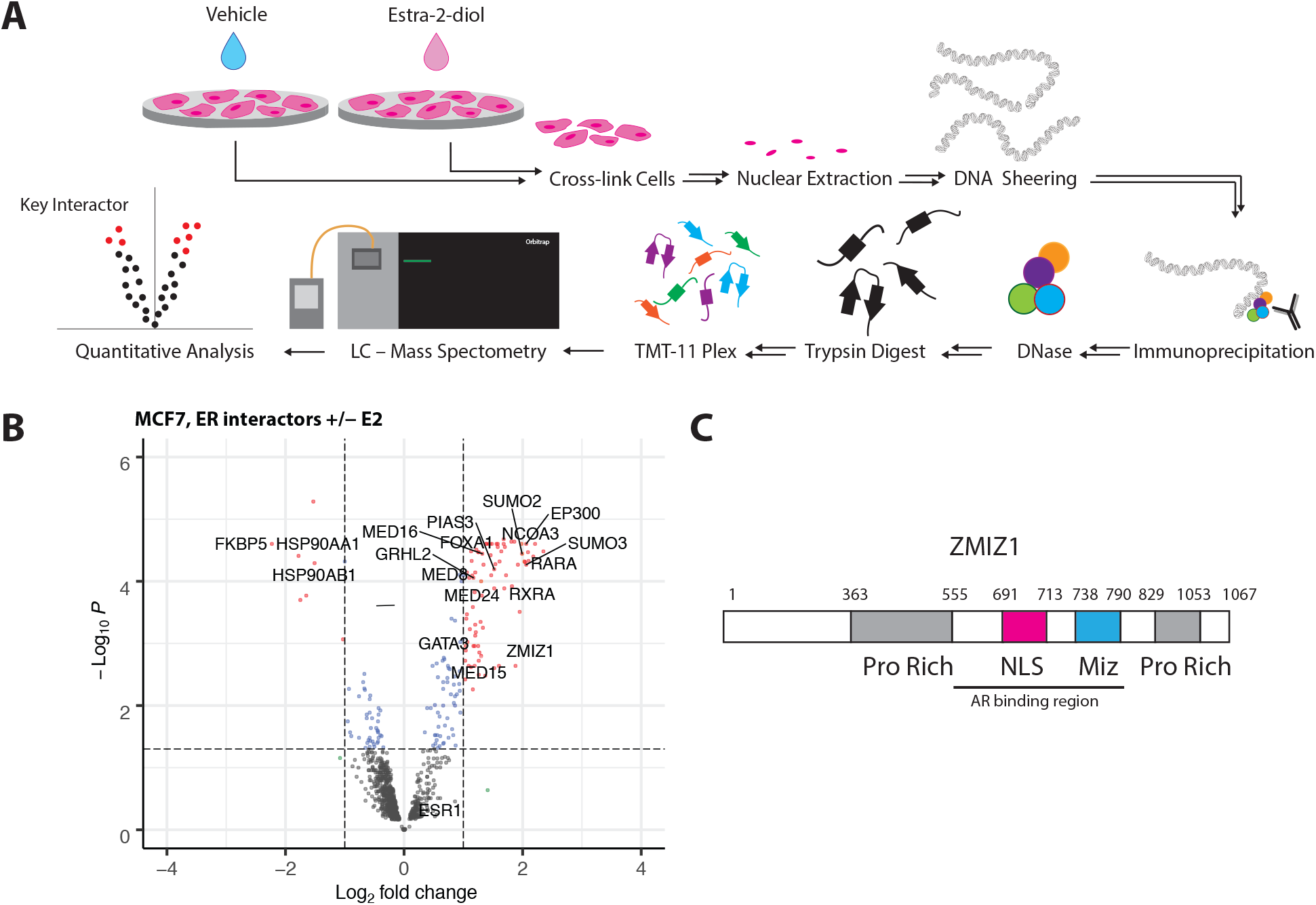
qPLEX-RIME of ER activation identifies novel co-factor ZMIZ1. **(A)** qPLEX-RIME enables the quantitative comparision of multiple conditions to idenfity key interactions between transcription factors on the chromatin. We stimulated ER+ breast cancer cell lines with estradiol and undertook a comparative analysis against an unstimulated control to identified the changes in the ER interacome on activation. **(B)** ER interacting proteins identified by qPLEX-RIME in MCF7. Top ranking differential-associated proteins detected with p < 0.05 and LogFC > 1 are highlighted in red. Gain of known co-factors GATA3 (Theodorou et al., 2013), RARA (Ross-Innes et al., 2010), EP300, GRHL2 (Holding et al., 2019) and NCOA3 were all detected along with the loss of HSP90 on binding of estradiol by the ER. Parts of the mediator complex (MED8, MED16, and MED24) along with pioneer factor FOXA1, were also detected. ZMIZ1 has not previously been reported to interact with the ER. *Caption continues on next page* **(C)** A schematic presentation of the ZMIZ1 protein. The AR binding region was previously identified by the interaction of a 556-790aa truncated mutant with a AR-GAL4 DBD fusion protein leading to activation of *β*-gal reporter gene. The C-terminal, proline-rich region of ZMIZ1, was identified as an intrinsic trans-activation domain (TAD) (Sharma et al., 2003).

Detected protein-ER interactions that were found to significantly change on stimulation with estradiol (p < 0.05) and had a two-fold change in protein intensity were reviewed for known biology. We found several previously identified ER co-factors including RARA, CBP, EP300, NRIP1 and GATA3. We also detected SUMO1-3 within the ER complex, most likely as a result of the covalent modification of the estrogen receptor or another protein within the ER chromatin-bound complex, in agreement with previous ChIP-seq experiments (Traboulsi et al., 2018). In addition, we also found putative novel protein partners of ER (Figure 1B).

The list of potential new ER co-factors includes ZMIZ1 (Figure 1C), which had previously been identified as a co-activator of the Androgen Receptor (AR) (Li et al., 2011; Sharma et al., 2003). However, a key result of that study is that the effect was AR specific in CV-1 cells. Transfection of ZMIZ1 into CV-1 cells was not able to enhance GR, PR, ER, and VDR-mediated transcription (Sharma et al., 2003). The AR specificity of the previous ZMIZ1 result has since been incorporated in the GeneCards and Uniprot databases (UniProt Consortium., 2019), and unpublished yeast data, discussed by (Sharma et al., 2003), supports the literature that there is no interaction between ZMIZ1 and non-AR nuclear receptors. Given that our qPLEX-RIME data shows a significant interaction between ER and ZMIZ1 in the breast cancer setting, and that we did not detect the presence of AR, we considered the breast cancer specific function of ZMIZ1 to be of key interest for follow-up studies.

### Proximity ligation assay validation of ER-ZMIZ1 interaction

We applied proximity ligation assay (PLA) to validate our qPLEX-RIME results. PLA signal is specific to proteins that are within 16nm (Trifilieff et al., 2011), and we successfully detected the proximity of ER and ZMIZ1 in both of the nuclei of two ER+ cell lines (MCF7 and T47D). Our analysis of the ER-ZMIZ1 interaction in the ER-cell line, MDA-MB-231, demonstrated no significant signal over our ER-IgG dual recognition PLA negative control, indicating the signal was specific to ER+ cell lines. To confirm the interaction of ER-ZMIZ1 was dependent on the protein-protein interaction, we treated the three cell lines with 100nM Fulvestrant, a Selective ER Degrader (SERD). On treatment, we measured a significant reduction (p<0.005 for MCF7 and T47D respectively) in the number of PLA signals in ER+ cell lines for both the ER-ER and ER-ZMIZ1 dual recognition PLA assays when compared to the vehicle control (Figure 2A–B). Additional PLA controls are shown in Supplementary Figure S2. We attempted to establish the interaction by co-IP (Figure S3) under native conditions, and successfully confirmed the interaction of ER and ZMIZ1 under native pull-down conditions in the T47D cell line, but not in the MCF7 or MDA-MB-231 cell lines. An explanation for the discrepancy in our results is that PLA detects proteins within the complex through space, and qPLEX-RIME uses cross-linking to both DNA and protein to capture transient interactions, while native Co-IP is more dependant on the affinities within the protein complex. Further, our T47D cells demonstrated higher expression of ER than MCF7, detected in both our ER-ER and in our ER-ZMIZ1 PLA signals, which provides a potential explanation of why the signal was detected in our T47D cell line over MCF7 cell line. Our results therefore confirmed the ER and ZMIZ1 are within the same protein complex in breast cancer cell lines, though the interaction maybe transient or indirect.

**Figure 2:**
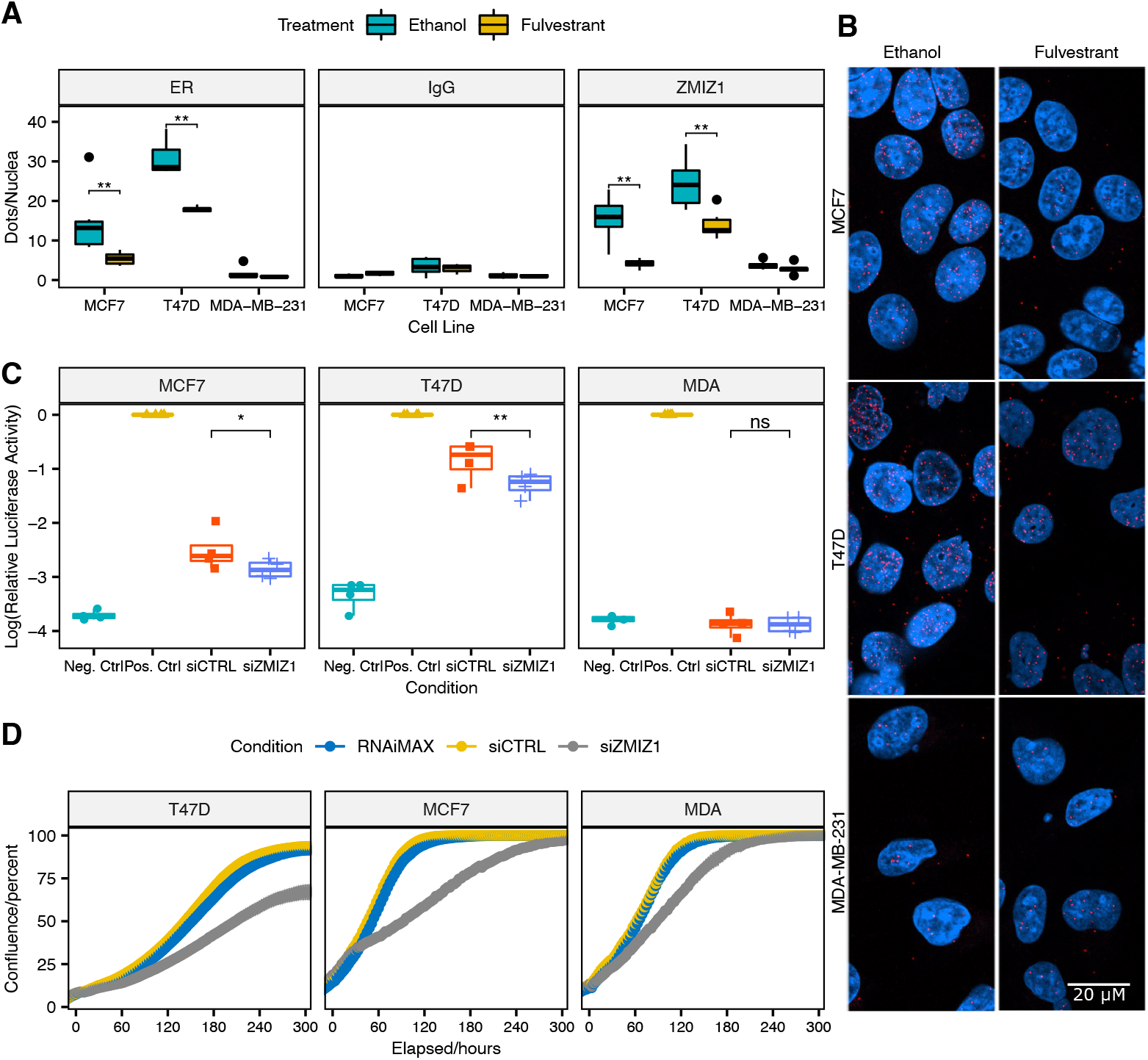
ZMIZ1 is within the ER transcriptional complex and ZMIZ1 knockdown reduces ER transcriptional activity and ER+ breast cancer cell line proliferation. **(A)** Cells were treated for 24 hours with vehicle (ethanol) or with 100nM Fulvestrant (a selective ER degrader) before analysis by PLA. *Caption continues on next page* Dual antibody recognition PLA between ER (Ab3575) and either ER (SC8002), IgG or ZMIZ1 demonstrated a significant reduction in both ER and the ER-ZMIZ1 interaction on addition of Fulvestrant in ER+ cell lines, indicating that ER and ZMIZ1 are within the same protein complex. No significant effect on the PLA signal was detected in the ER-IgG negative control, or in the ER-cell line MDA-MB-231. (** Wilcoxon signed-rank test, p-value < 0.005). The error bars indicate standard deviation (SD). **(B)** Illustrative images of the ER-ZMIZ1 PLA, taken at 630x magnification. DAPI stained cell nuclei are shown in blue and PLA signals are shown as red dots. **(C)** Lucierase Assay monitoring ER activity. ER activity in MCF7, T47D and MDA-MB-231 cell lines was monitored by Cignal ERE Reporter Assay. Both MCF7 and T47D cells showed a significant reduction in ER activity in response to ZMIZ1 knockdown. As expected, MDA-MB-231 cells showed no activity in all conditions but the positive, constitutively active, control. (*/** one-sided t-test, p-value < 0.05/0.005.) **(D)** Knockdown of ZMIZ1 in three cell lines, T47D, MCF7 and MDA-MB-231 all showed reduced proliferation. The effect was largest in the ER-positive cell lines, T47D and MCF7. We inferred the increased response to ZMIZ1 knockdown to imply that as ER signalling is the main driver in these two cell lines, that ZMIZ1 expression levers had the greatest effect. In contrast, the response in MDA-MB-231, a triple-negative breast cancer model, implies that ZMIZ1 has additional functionality independent of the estrogen receptor transcriptional complex, but this has a lesser effect on cell growth than its role in ER positive breast cancer.

### ZMIZ1 knockdown reduces ER transcriptional activity

We hypothesised that as our breast cancer model was able to support the ER-ZMIZ1 interaction, that ZMIZ1 may have a transcriptional role in the ER complex.

To determine if ZMIZ1’s role within the ER complex regulated transcriptional activity, we monitored the changes in ER activity on knockdown of ZMIZ1 with siRNA using a Cignal Reporter Assay. We used two models of ER+ breast cancer, MCF7 and T47D, along with MDA-MB-231, a model of Triple Negative Breast Cancer. Estrogen receptor activity was measured by a luciferase activity assay relative to a constitutive active positive control. Renilla luciferase activity was used to control for transaction efficiency.

Both MCF7 and T47D (Figure 2C) cell lines showed significantly reduced ER activity (p < 0.044 and p < 0.0045 respectively, paired t-test, one-tailed, n = 4) with siZMIZ1 compared to the control (siCTRL). As expected, MDA-MB-231 showed no detectable ER activity, and therefore no significant difference between the siZMIZ1 and siCTRL conditions or between the experimental conditions and the negative control. These results confirm that ZMIZ1 has a co-activator role at ER regulated genes, and the effect is dependant on ER expression.

### ZMIZ1 knockdown delays response to E2 in ER regulated cell-cycle related genes

Since knockdown of ZMIZ1 leads to a significant reduction in ER transcriptional activity, we hypothesised that knockdown of ZMIZ1 would also result in a reduced response to E2 at downstream targets of the ER.

To test our hypothesis, we set up a paired RNA-seq experiment treating MCF7 cells with either siZMIZ1 or siCTRL. In both cases, we undertook four isogenic replicates, measuring transcriptional levels at 3, 6, 12 and 24 hours after stimulation with estradiol. The reduction in expression of ZMIZ1 by siRNA knockdown was found significant at all time points compared to the control (adj-p value < 0.01, n=4, Figure S1), No significant change in the expression of the ER was detected.

Between the siZMIZ1 and control condition, the largest number of differentially expressed genes occurred at 6 hours after stimulation with E2 (30 hours after initial knockdown). In contrast, by 24 hours (48 hours after knockdown of ZMIZ1) only 14 genes were detected as differentially expressed.

Analysis of the 6 hour time-point (siZMIZ1 vs siCTRL) by Gene Set Enrichment Analyis (GSEA) (Subramanian et al., 2005) for enrichment of ER responsive genes gave conflicting results. We identified three published gene sets relevant to the cell culture models used: STEIN_ESR1_TARGETS and `BHAT_ESR1_TARGETS_NOT_VIA_AKT1_UP` described the genes regulated by the ER in the MCF7 cell line, while `WILLIAMS_ESR1_TARGETS_UP` reflected genes activated on the stimulation of the ER with estradiol in the T47D cell line. GSEA analysis of the differential expression at 6 hours of stimulation with E2 in MCF7 for `STEIN_ESRl_TARGETS` gave a non-significant reduction (p = 0.07), `BHAT_ESR1_TARGETS_NŨT_VIA_AKTl_UP` gave a significant increase (p = 0.003), and `WILLIAMS_ESR1_TARGETS_UP` showed a significant reduction in ER transcriptional response (p = 0.047). Overall overlap of all 3 gene sets is low with 3 genes in common. The STEIN_ESR1_TARGETS gene set has 25% overlap with the other two sets, while the `WILLIAMS_ESR1_ TARGETS_UP` gene set and `BHAT_ESR1_TARGETS_NOT_VIA_AKTl_UP` have 46% and 9% overlap respectively with the rest of the gene sets. Given the ER’s role in regulating cell-cycle (JavanMoghadam et al., 2016), we undertook GSEA against three MSigDB gene sets (Subramanian et al., 2005) focused on cell-cycle: `GO_CELL_CYCLE, KEGG_CELL_CYCLE`, and `REACTOME_CELL_CYCLE`. All three gene sets gave significant results (p = 2.3 × 10^-6^,0.0001, and 3.5 ×10^-7^ respectively, Figure S6).

On the basis of these results, we hypothesised that ZMIZ1 regulated a subset of ER-regulated genes focused on cell-cycle. The varied results for GSEA of ER specific sets could then be explained as a result in variation in genes represented in each of MSigDB gene sets and a lack of specificity to ER-ZMIZ1 signalling axis. To test our hypothesis, we undertook GSEA against the intersection of the `REACTOME_CELL_CYCLE` and the previously tested ER specific gene sets (p = 0.0006). Repeating the analysis in T47D cell gave p = 0.005 (Figure 3A).

**Figure 3:**
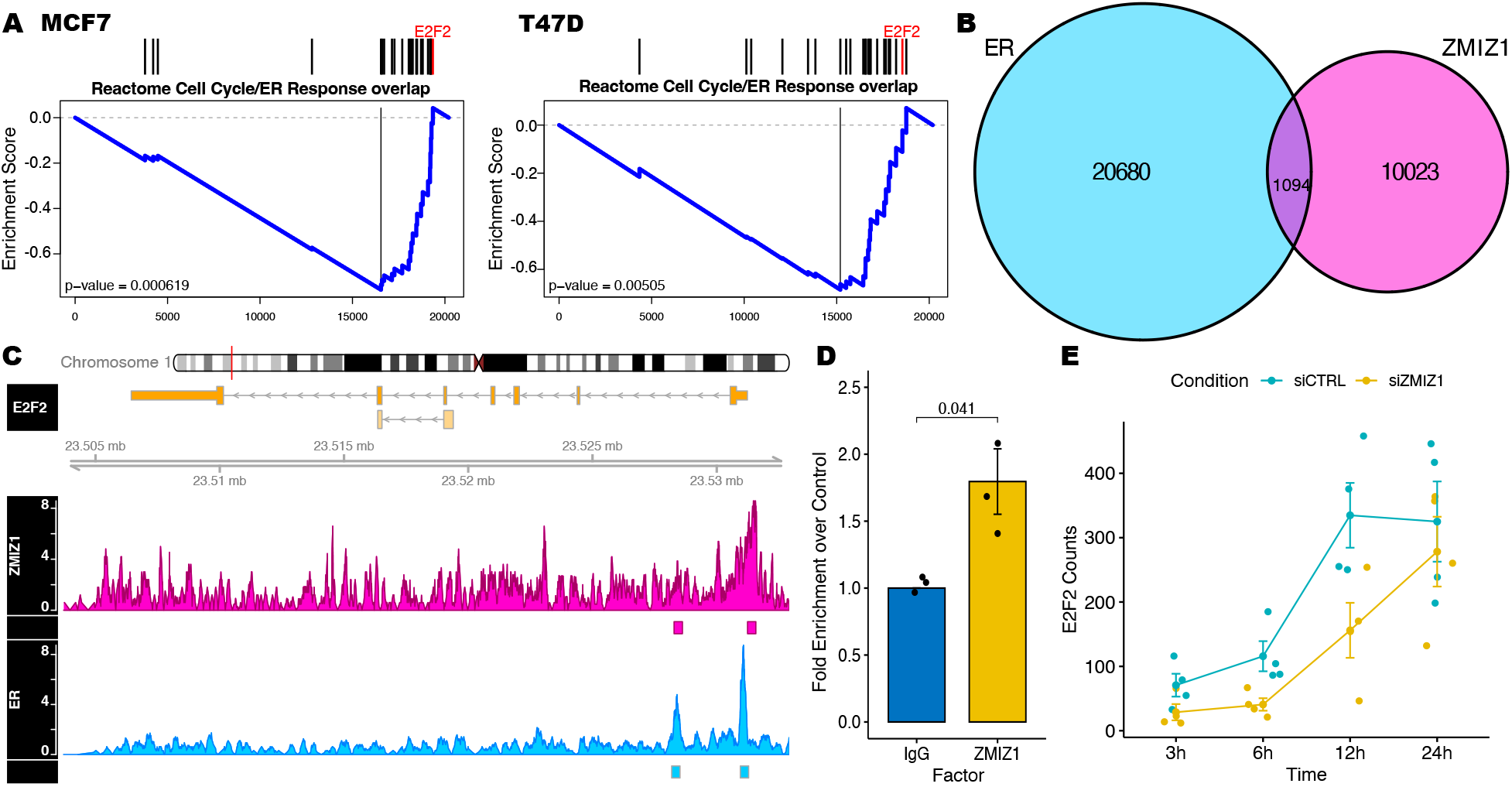
ZMIZ1 knockdown leads to a delay in the transcriptional response of estrogen regulated cell-cycle genes. ER and ZMIZ1 co-localise at E2F2 promoter. **(A)** GSEA of RNA-seq data generated 6 hours after stimulation of cells by estrogen found significant depletion in our gene set created from the intersection of cell-cycle and estrogen response genes in cells with ZMIZ1 knocked down. **(B)** Overlap of ZMIZ1 binding in K562 cells (ENCFF881DAT) with ER binding in MCF7 (ENCFF138XTJ) downloaded from ENCODE showed that 10% of ZMIZ1 binding sizes overlapped with ER. **(C)** Analysis of sites found at cell-cycle genes co-occupied by ER and ZMIZ1 identified co-binding within the promoter of E2F2, a gene also found within our gene set. Coverage maps for ZMIZ1 and ER (ENCFF042TOP, ENCFF237WTX) showed a clear peak in the ChIP-seq signal for both factors at the promoter of the E2F2 gene. A second peak within the first intron was also annotated in both data ENCODE sets, but the peak at the annotated site was not visually distinguishable from background in the ZMIZ1. **(D)** ChIP-qPCR validation of the ZMIZ1 binding site identified in the ENCODE ChIP-seq data confirmed significant enrichment of E2F2 promoter DNA in the ZMIZ1 pull-down over IgG control in MCF7 cells. **(E)** Reanalysis of the E2F2 transcript within our RNA-seq data showed reduced expression at 3-12 hours.

### ZMIZ1 and ER co-bind and co-regulate cell-cycle regulator E2F2

As ZMIZ1 and ER are both transcription factors, we aimed to establish if the proteins co-bind DNA elements that regulate cell-cycle. We surmised any sites we identified could indicate a potential cell-cycle regulatory mechanism ER-ZMIZ1 interaction. For ZMIZ1 binding we used the ENCODE data reported in K562 cells, as no data for MCF7 cells was available, and for ER-binding we used data as reported by ENCODE in MCF7 cells (Luo et al., 2020). Overlaying these data demonstrated nearly 10% of ZMIZ1, 1094 sites in total, were found to co-bind ER (Figure 3C) in support of our PLA results. Comparison of the ER–ZMIZ1 co-bound sites with our GSEA gene set identified E2F2 as being within both data sets. ENCODE reported two binding-sites within the E2F2 gene loci for both ER and ZMIZ1, and both sites overlapped. Visual inspection of the ENCODE data showed clear binding of ER and ZMIZ1 at E2F2 promoter region. While the second binding site within the E2F2 intron was clearly bound by ER, the intronic ZMIZ1 ChIP-seq peak was at similar levels to background noise (Figure 3C). To address concerns that the ENCODE ChIP-seq data was generated in K562 cells, we validated the ZMIZ1 binding of the E2F2 promoter in MCF7 cell line by ChIP-qPCR confirming our ENCODE ZMIZ1 ChIP-seq result was generalisable to our breast cancer model (Figure 3D). Reanalysis of the RNA-seq data showed a reduction in E2F2 expression, with a maximum log_2_-fold change of −1.6 at 3h, and a log_2_-fold change of −1.4 between 6–12 hours (Figure 3E). Suggesting, that our knockdown of ZMIZ1 resulted in a delayed increase in the expression of the E2F2 transcriptional factor in response to estrogen due to the reduction in ER-ZMIZ1 interaction at the E2F2 promoter.

### ZMIZ1 knockdown reduces proliferation of breast cancer cell lines

On the basis of these results, we tested the hypothesis that knockdown of the ZMIZ1 protein would result in reduced proliferation of ER-positive cancer cell line models.

Analysis of two ER-positive (MCF7 and T47D) and one triple negative breast cancer model (MDA-MB-231) showed that knockdown of ZMIZ1 reduced cell proliferation in all three cell lines (Figure 2D). The effect was greatest in the ER positive cell lines, T47D and MCF7, with T47D growth reaching a maximum at a reduced confluence compared to the control ex-periments. In contrast, the magnitude of the reduction in growth was more modest in the MDA-MB-231 cell line and the rate of recovery was much higher. The susceptibility of T47D and MCF7 to ZMIZ1 knockdown is likely due to the specific reliance of these cells on ER for cell growth, while in the MDA-MB-231 cell line ZMIZ1 likely regulates a second ER-independent pathway.

### High ZMIZ1 expression correlates with low survival in ER+ patients

To explore if the role of ZMIZ1 activity held clinical importance we investigated if ZMIZ1 expression was a predictor of patient survival in the context of both ER+ and ER-tumours using KMplot (Lanczky et al., 2016).

Stratifying ER+ patients by median ZMIZ1 expression showed that ER+ patients with higher levels of ZMIZ1 expression had significantly poorer out-come (p = 0.0023, Logrank test).

In contrast, the same analysis of ER-negative patients demonstrated no significant difference in patient outcome (p = 0.49, Logrank test, Figure 4A). Survival differences were also confirmed in METABRIC and TCGA ER+ breast cancer cohorts (Figure S5).

**Figure 4:**
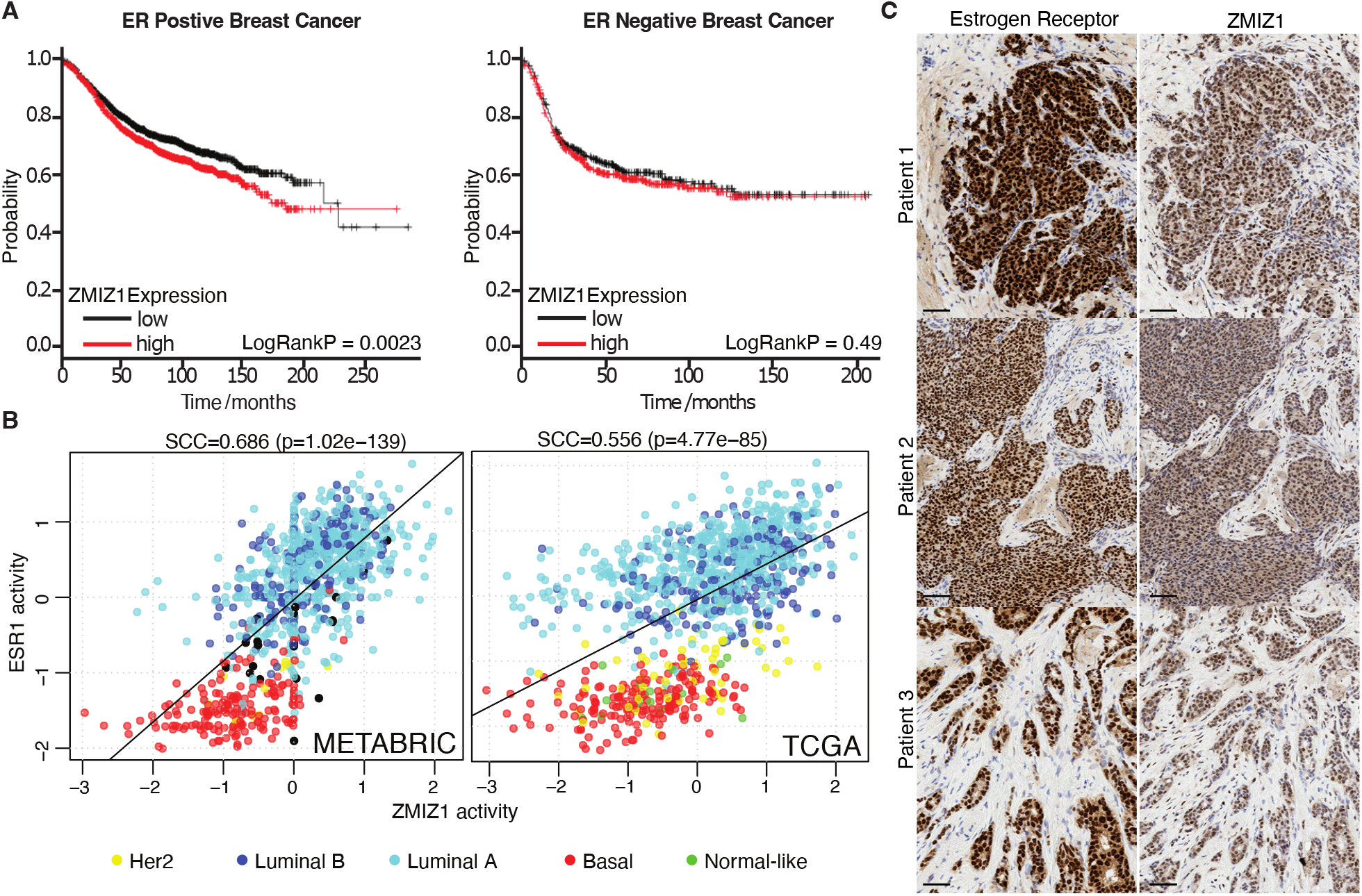
High ZMIZ1 expression correlates with low survival in ER+ patients, ZMIZ1 and ER transcriptional activities correlate in patient samples, and ZMIZ1 and ER are co-localised in patient samples. **(A)** Stratifying patients of ER+ (left) breast cancer based on ZMIZ1 expression shows that high levels of ZMIZ1 results in a significant reduction in patient survival. The effect is not seen with ER -ve (right) breast cancer, implying an ER+ specific function for ZMIZ1 in breast cancer. **(B)** Transcriptional activity of ESR1 and ZMIZ1, as measured by the VIPER algorithm, correlate in both the METABRIC and TCGA cohort thus confirming both transcription factors are transcriptionally active within ER+ tumours and potentially function together in ER+ tumours. **(C)** Consecutive slides derived from biopsy samples of three patients with ER+ breast cancer were stained by IHC for both the Estrogen Receptor-alpha and ZMIZ1. In all three patients, the localisation of staining for the proteins was specific to the tumour cells and the nucleus. In contrast, the stromal cells show little or no expression of either protein. Scale = 50 *μ*m.

These results were consistent with the ZMIZ1 functioning as a co-factor of the ER in ER-positive breast cancer and that ZMIZ1 played a role in disease progression in ER-positive breast cancer.

### ZMIZ1 and ER activities correlate in patient samples

If ZMIZ1 is a key component of the ER complex, we would expect its activity to be higher in ER-positive than in ER-negative breast cancer. To test this hypothesis, we used two large gene expression collections, TCGA (Weinstein et al., 2013) and METABRIC (Curtis et al., 2012), to assess co-expression of ER and ZMIZ1 in a patient setting. In both datasets, ZMIZ1 had significantly increased expression in luminal over basal sub-types (TCGA: p=1.34× 10^-23^, Wilcoxon test, n=962; METABRIC: p=1.18× 10^-10^, Wilcoxon text, n=961).

To address if both ZMIZ1 and ER proteins were transcriptionally active within patient samples, we then generated regulatory network models for both the TCGA and METABRIC data sets using ARACNe-AP (Lachmann et al., 2016). Using these networks, we applied the VIPER algorithm (Alvarez et al., 2016) to calculate the activity of ER and ZMIZ1 proteins in both patient sample datasets (Figure 4B). For both TCGA and METABRIC, we saw a significant correlation (p=4.77 × 10^-85^ and p=1.02× 10^-139^ respectively) between the activity of the two transcription factors, with greatest activity of both TF networks in the luminal sub-type.

### Immunohistochemistry of ZMIZ1 and ER demonstrates co-localisation in patient samples

To further validate the link between ER and ZMIZ1, we checked if ZMIZ1 expression was co-localised with that of the ER in clinical material. Visual inspection of ER-positive of samples breast cancer tumours from all three patients analysed showed strong nuclear staining of both proteins in adjacent sections. Comparison of the localisation of staining between ER and ZMIZ1 demonstrated that both proteins were found within the nucleus of epithelial cells. Neither ER, nor ZMIZ1, was not detected in the infiltrating cells. Further, the distribution of ER and ZMIZ1 staining correlated, suggesting the ER and ZMIZ1 are expressed within the same cells of the patient tumours (Figure 4C).

## 3. Discussion

Previous evidence supported the proposal that ZMIZ1 was a specific co-activator of the androgen receptor (Sharma et al., 2003). Results in the cited study demonstrated that AR, ER, and other steroid hormone receptors could all activate transcription when presented with their respective ligands in the cell line model used. However, only the activity of AR was linked to the levels of ZMIZ1 within the cell. On detection of ZMIZ1 within the ER complex, we therefore hypothesised that the absence of evidence for an ER-ZMIZ1 interaction until this point may be a result of the cell line model used in that study.

Undertaking analysis in both the MCF7 and T47D cell lines (ER-positive breast cancer models) demonstrated a significant change in ER-mediated transcriptional activity on ZMIZ1 knockdown. We therefore present evidence that ZMIZ1 has a role in modulating ER activated transcription that has not previously been seen in other models. Further, we were able to show that within patient samples the ZMIZ1 and ER expression correlated, and that the proteins are co-localised within the nuclei of tumour cells.

In this study, we demonstrate that ZMIZ1 is able to interact with the ER complex and that loss of ZMIZ1 results in lower transcriptional activity. In particular, reducing the proliferate cycle-cell response to estrogen via a novel ER-ZMIZ1-E2F2 signalling axis. Previously it has been shown that ZMIZ1 has a very strong trans-activation domain (TAD) (Sharma et al., 2003) and our study is consistent with this mechanism, suggesting that ZMIZ1 is a co-activator within the ER complex. Alternatively, ZMIZ1 may activate the ER by promoting the SUMOylation of either the ER or its co-factors. A similar role has been seen for the AR, increasing SUMOylation by about 40% (Sharma et al., 2003), and our own qPLEX-RIME analysis of the ER shows an increase in the identified SUMO protein modifications on stimulation estradiol. However, the mechanism by which ZMIZ1 can promote SUMOylation remains unclear.

All nuclear receptors orchestrate their response on activation by interactions with large numbers of co-regulators (Rosenfeld & Glass, 2001). Loss or gain of function mutations that affect these co-factors may then result in a selective advantage for the tumour, or even lead to resistance to treatment (Wu et al., 2018). In this context, ZMIZ1’s role in mediating AR driven cell growth (Sharma et al., 2003) has led to suggestions that the protein holds an interesting opportunity for targeting prostate cancer disease progression during androgen deprivation therapy (Li et al., 2011). In breast cancer, the ER’s role in driving cell growth and proliferation is a key factor in patient survival and levels of ZMIZ1 are a significant predictor of patient outcome. These features of ZMIZ1 suggest similar potential for targeting the protein as a therapeutic opportunity limiting ER-positive breast cancer growth. This hypothesis is further supported that our analysis of the transcriptome after knockdown of ZMIZ1 showed a specific and significant reduction in estrogen-mediated cell-cycle gene activation.

## 4. Conclusion

In conclusion, we have shown that loss of ZMIZ1 reduces the activity of ER-activated genes and slows cell growth in ER+ breast cancer cell lines. On the molecular level, we have demonstrated ER and ZMIZ1 co-localise within the nucleus and that loss of ZMIZ1 results in a delayed expression of estrogen-responsive cell cycle genes. This result is supported by the conformation of the ER-ZMIZ1 interaction at the E2F2 promoter, itself a cell cycle regulator and one of the most down regulated genes in response to ZMIZ1 loss, thereby providing a mechanism for the effect of ZMIZ1 knockdown on E2-mediated cell cycle. Finally, we demonstrate the clinical relevance of this interaction by confirming that ZMIZ1 is expressed within ER+ patient tumours through the analysis of patient biopsies, and by showing that increased ZMIZ1 expression correlates with worse outcomes in ER+ patients.

We believe these results highlight the importance of further studies into the molecular basis by which ZMIZ1 regulates both ER, and AR, mediated transcription; as they may provide new insight into the biological role of ZMIZ1 in hormone driven cancers along with novel opportunities for therapeutic intervention specific to tumour growth.

## 5. Materials and Methods

### Cell lines and general cell culture

MCF7, T47D (ER positive cell lines) and MDA-MB-231 (ER negative cell lines) were obtained from ATCC. All cells were routinely cultured in DMEM with high glucose and pyruvate (Gibco #41966-029) supplemented with 10% FBS. Cells were confirmed to be mycoplasma free using the MycoStrip test supplied by InvivoGen. Cell line authentication was confirmed by short tandem repeat genetic profiling carried out by Eurofins. All cell lines were used at passages less than 35 for all experimental work described below.

### qPLEX-RIME

qPLEX-RIME samples were prepared as previously reported (Papachristou et al., 2018). Cells were grown in estrogen-free culture as previously described (Guertin et al., 2018). In summary, cells were cultured for four days in phenol red-free Dulbecco’s Modified Eagle’s Medium (Glibco) with 10% charcoal stripped Fetal Bovine Serum (FBS) and glutamine, cells were washed with PBS and media was changed daily to remove residual estrogen. Cells were cross-linked for qPLEX-RIME 45 minutes after addition of 100 nM estradiol (E2) (Sigma-Aldrich) prepared in ethanol or ethanol (control).

### Proximity Ligation Assay (PLA)

MCF7, T47D and MDA-MB-231 were seeded into 8 well chamber slides in full media. After 48 hours (around 50% confluent), half of the wells were treated with 100nM of Fulvestrant (Cayman #10011269) and the other half were treated with an equivalent amount of vehicle control (Ethanol) for 24 hours. After 24 hours of treatment, cells were rinsed in PBS and fixed with ice-cold methanol. Prior to performing the PLA assay, methanol was removed and samples rinsed in PBS. The PLA assay (Sigma Aldrich) was then performed according to the manufacturer’s instructions. To assess for the close proximity of ER and ZMIZ1, anti-ER antibody supplied by Abcam (Ab3575) diluted at 1:400 and anti ZMIZ1 supplied by Santa Cruz (SC-376825) diluted at 1:250 were used together. Dual antibody recognition of ER using anti-ER antibody from Abcam (Ab3575) at 1:400 dilution and anti-ER antibody from Santa Cruz (SC8002) at 1:250 was used as a positive control. An isotype anti-IgG antibody from Cell Signaling (27295) diluted to 1:250 was used, together with the anti ER antibody from Santa Cruz antibody at 1:250, as a negative control. Following DUOLINK staining, the samples were imaged using a LSM780 confocal microscope at x630mag. 2 replicate wells were prepared for each sample condition and 4 different fields of view were imaged from each sample well. Images were then analysed using Image J software. The number of PLA signals (fluorescent dots) to number of cells per image was assessed using the ‘Analyse particle’ function in the Image J software. All statistical analysis was carried out using Wilcoxon signed-rank test.

### ERE activity assay

Four biological replicates of MCF7, T47D and MDA-MB-231 were assayed using Cignal ERE Reporter Assay Kits (Qiagen) according to the manufacturer’s protocol. Estrogen-free culture of all three cell lines was established as previously reported for MCF7s (Guertin et al., 2018). SMARTpool ZMIZ1 (L-007034-00)and control siRNA (D-001810-10, Dharmacon) were transfected with Lipofectamine RNAiMAX Transfection Reagent (Thermo Fisher Scientific) following the manufacturer’s protocol.

### Cell growth assay

MCF7, T47D and MDA-MB-231 cell growth was undertaken in complete-DMEM with 10% FBS and glutamine, monitored by incucyte. siRNA knock down was undertaken with SMARTpool on-target siZMIZ1 along with the matched siCTRL (Dharmacon) and was transfected with Lipofectamine RNAiMAX reagent (Thermo Fisher Scientific) according to the manufacture’s protocol. Cells were reverse transfected, and media was refreshed after 24 hours to minimise the toxicity of the transfection reagents. Knockdown was confirmed by RT-qPCR.

### RNA-sequencing and GSEA

MCF7, T47D and MDA-MB-231 cell growth was undertaken in phenol red-free DMEM +10% Charcoal Stripped FBS. siRNA knockdown was undertaken with SMARTpool on-target siZMIZ1 along with the matched siCTRL (Dharmacon) and was transfected with Lipofectamine RNAiMAX reagent (Thermo Fisher Scientific) according to the manufacture’s protocol. 24 hours after transfection, the cell culture media was refreshed, to minimise any toxic affect of the transfection reagent, and cells were either stimulated with 100nM E2 or ethanol as a control. After 3, 6, 12 and 24 hour stimulation with either E2 or ethanol, RNA was collected for 4 biological replicates within each cell line and libraries were prepared by TruSeq mRNA Library Prep (Illumina). The libraries were sequenced on the Illumina HiSeq 4000 platform. Reads were aligned with HiSat2 to hg19 and count matrix established with Htseq-Count v2.1.0. Differential analysis of expression was undertaken with the R-package DESeq2 (Love et al., 2014), and GSEA(Mootha et al., 2003; Subramanian et al., 2005) was undertaken using the implementation in the R-package VULCAN (Holding et al., 2019).

### ChIP-qPCR

ChIP-qPCR was undertaken according to the published methods(Glont et al., 2019) with some modifications. A total of 40 million MCF7 cells were used as starting material per each ChIP reaction and three isogenic replicates were prepared from independent cell culture. With the cell culture medium being discarded, MCF7 cells were crosslinked for 10 minutes by 1% formaldehyde solution (Thermo Fisher Scientific, 28908) in PBS at ambient temperature followed by quenching with 0.125 M glycine for 5 minutes. After washing with cold PBS twice, fixed cells were collected with cold PBS supplemented with 1X cOmplete protease inhibitor cocktail (Sigma Aldrich, 11873580001). The cell pellets were resuspended and incubated at 4°C in 1 mL Lysis Buffer 1 for 10 minutes and in 1 mL Lysis Buffer 2 for 5 minutes sequentially. After each incubation step, samples were centrifuged at 5000 g for 5 minutes to recover the cell/ nuclei pellets. The pellets were resuspended in 300 *μ*L Lysis Buffer 3 and sonicated by a Bioruptor Pico (Diagenode), resulting in chromatin fragments of around 300 to 500 bps. Buffer were prepared as previously reported(Glont et al., 2019). After sonication, the cell lysates were supplemented with Triton X-100 (Sigma Aldrich) at a final concentration of 1% and were centrifuged at 13500 RMP for 10 minutes at 4°C to remove debries. 30 *μ*L of supernatant was saved as input for sub-sequent qPCR analysis. The rest of the supernatant was mixed with prior prepared antibody-conjugated protein A/G beads (Thermo Fisher Scientific, 88802) and was incubated at 4°C overnight. 5 *μ*g of anti-ZMIZ1 antibody (Abnova, PAB4820) or normal rabbit IgG (Cell Singaling Technology, 2729) was used for each ChIP reaction. After overnight incubation, the beads were washed with RIPA buffer (50 mM HEPES pH 7.6, 1mM EDTA, 1% NP40, 0.5M lithium chloride, 0.7% sodium deoxycholate) six times in a cold room, followed by a rapid wash with TE buffer (pH 7.0, Invitrogen AM9861). The crosslinks of both ChIP samples and input were reversed in 200 *μ*L ChIP Elution Buffer (50 mM Tris-HCl pH 8.0, 10 mM EDTA, 1% SDS) at 65 °C overnight. Samples were subject to RNAse A (Invitrogen, AM2271) digestion at 37 °C for 30 minutes and Proteinase K (Invitrogen, 25530049) digestion at 65 ^°^C for 2 hours after reverse crosslinking. ChIP DNA was recovered by a QIAquick PCR purification kit (Qiagen, 28104) for subsequent qPCR analysis.

Primers probing the promoter of E2F2 (Forward 5’-CAGCTTGGGAGA GTAGAAGAAC, Reverse 5’-CCAAGGTCATACAGAGAGATTCC) were designed by IDT PrimerQuest Tool. qPCR reactions were set up with Fast SYBR Green Master Mix (Thermo Fisher Scientific, 4385612) per the users’ manual, where 10 *μ*L master mix, 1 *μ*L primers (250 nM), 1 *μ*L ChIP DNA and 8 *μ*L nuclease-free water were mixed per reaction. Technical triplicate was prepared for each reaction. All the qPCR assays were conducted in an Admiral QuantStudio 3 Real-Time PCR System (Applied BioSystems) under the default fast, comparative CT thermo profile. Statistical analysis was performed using paired, one-tailed t-test.

### Survival analysis

Initial survival analyses with undertaken using KMplot.(Győrffy, 2021) Patients were stratified on ZMIZ1 expression with median cut-off selected. ER+ status was established from array data. These settings on 11 August 2022 gave p-value < 0.001 for ER+ (n=5526) patients, and not significant for ER- (n=2009). METABRIC and TCGA survival data was downloaded from cBioPortal (Cerami et al., 2012), analysis was undertaken in R, the most suitable cut-off for expression was calculated using “survminer” package and data was fit using the ‘survival’ package.

### Network Analysis of ER and ZMIZ1 activity

Expression data from TCGA and METABRIC was analysed using VIPER (Alvarez et al., 2016) and ARACNe-derived networks (Lachmann et al., 2016) from the R-package ‘aracne.networks’ in Bioconductor. Statistical analysis and plotting of results was undertaken in R.

### Immunohistochemistry

Antibody specificity was confirmed on formalin fixed cell pellets of MCF7 cells (Figure S7). Patient tissue samples were run on Leica’s Polymer Refine Detection System (DS9800) using their standard template on the automated Bond-III platform. MCF7 cells were cultured in complete-DMEM with 10% FBS and glutamine. Post-siRNA knockdown of ZMIZ1 were used as negative control to ensure antibody specific. De-waxing and re-hydration prior to IHC automated on the Leica ST5020, along with the post-IHC dehydration and clearing. Sections were mounted using Leica’s coverslipper CV5030. The specific antibody targeting ZMIZ1 was purchased from R&D Systems (AF8107) and used at a concentration of 2 *μ*g/ml (1:250 dilution). The sodium citrate pre-treatment was run at 100 °C. The secondary (post primary) was rabbit anti-sheep from Jackson ImmunoResearch (r313-005-003), diluted 1:500. DAB Enhancer is included as an ancillary reagent (Leica, AR9432).

Patient tissue samples were processed as described for the MCF7 fixed cell pellets. Estrogen Receptor antibody was purchased from Novacastra (NCL-ER-6F11/2) and samples processed previously described (Bruna et al., 2016). All samples are from the PIONEER trial, REC number 17/NE/0113.

## 6. Data and code availability

All RNA-seq data have been deposited in the GEO database with the accession number GSE133381. The code and data to generate the figures in this paper are available as an R-package from https://github.com/andrewholding/ZMIZ1.

## 7. Acknowledgments

We would like to acknowledge the support of The University of York, York Biomedical Research Institute, The University of Cambridge and Cancer Research UK.

This work was funded by CRUK core grant C14303/A17197 and A19274 (to FM) and supported by The Alan Turing Institute under the EPSRC grant EP/N510129/129/1 as a Turing Fellowship, BBSRC grant BB/V000071/1, and Royal Society Research Grant RGS/R2/202120 to ANH.

We would like to acknowledge support of the Imaging and Cytometry group in the Technology Facilities at The University of York and the contribution from the CRUK Genomics, Proteomics, Histopathology/ISH, and Bioinformatics core facilities in supporting this work.

## 8. Author Contributions

WZ,JS, AG, RB, AEC, LB, SFR and ANH undertook the experimentation. SK provided patient material. AG and SK generated ZMIZ1 stained images of patient samples. JS. AG, RB, KK, AS, ME, FMG, SFR and ANH analysed the data. MG analysed the ZMIZ1 stained images of patient samples. FMG undertook network analysis of TGCA and METABRIC datasets. AG, KSB and ANH designed the experiments. WZ, AG, SFR, ANH and FM wrote the manuscript. All authors were involved in the proofing and editing of drafts.

