## Supplemental Figures for "In Breast Cancer ZMIZ1 Co-Regulates E2F2 as Part of the Estrogen Receptor-Mediated Cell-Cycle Response"

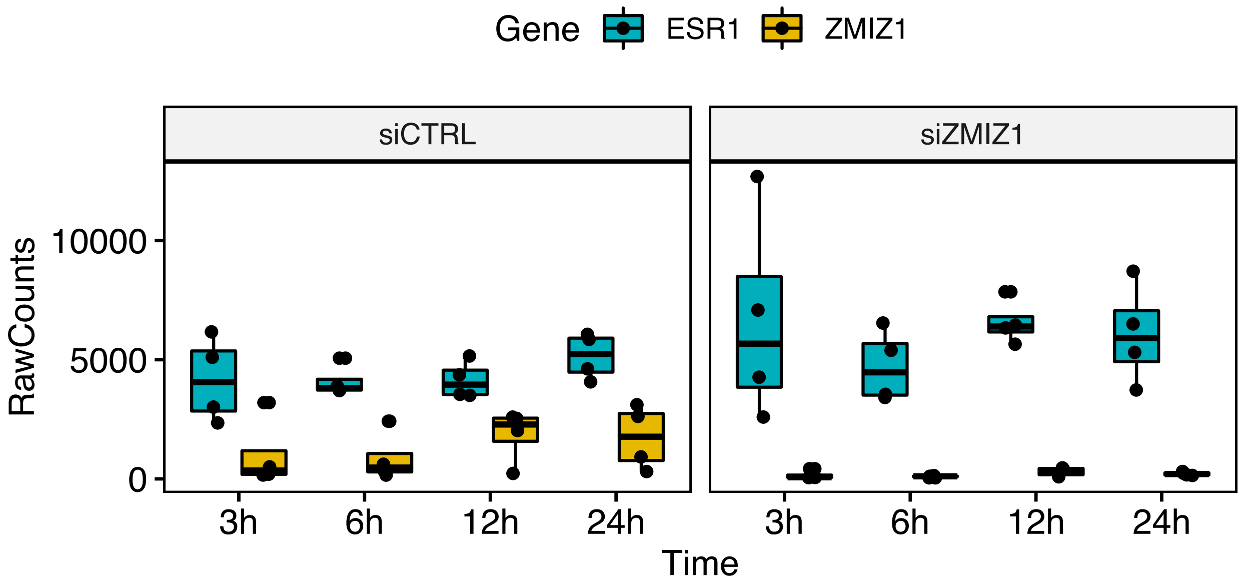


Figure S1: **Knockdown of ZMIZ1 by siRNA in MCF7 cells was detected at all time points**. Plot shows the number of raw reads aligned to ZMIZ1 and ESR1 in sample at each time point. Analysis of ZMIZ1 expression at all time points by DeSEQ2 found the transcript significantly reduced in expression ($p < 0.01$). ER expression was not found to change significantly on ZMIZ1 knockdown at any time point.

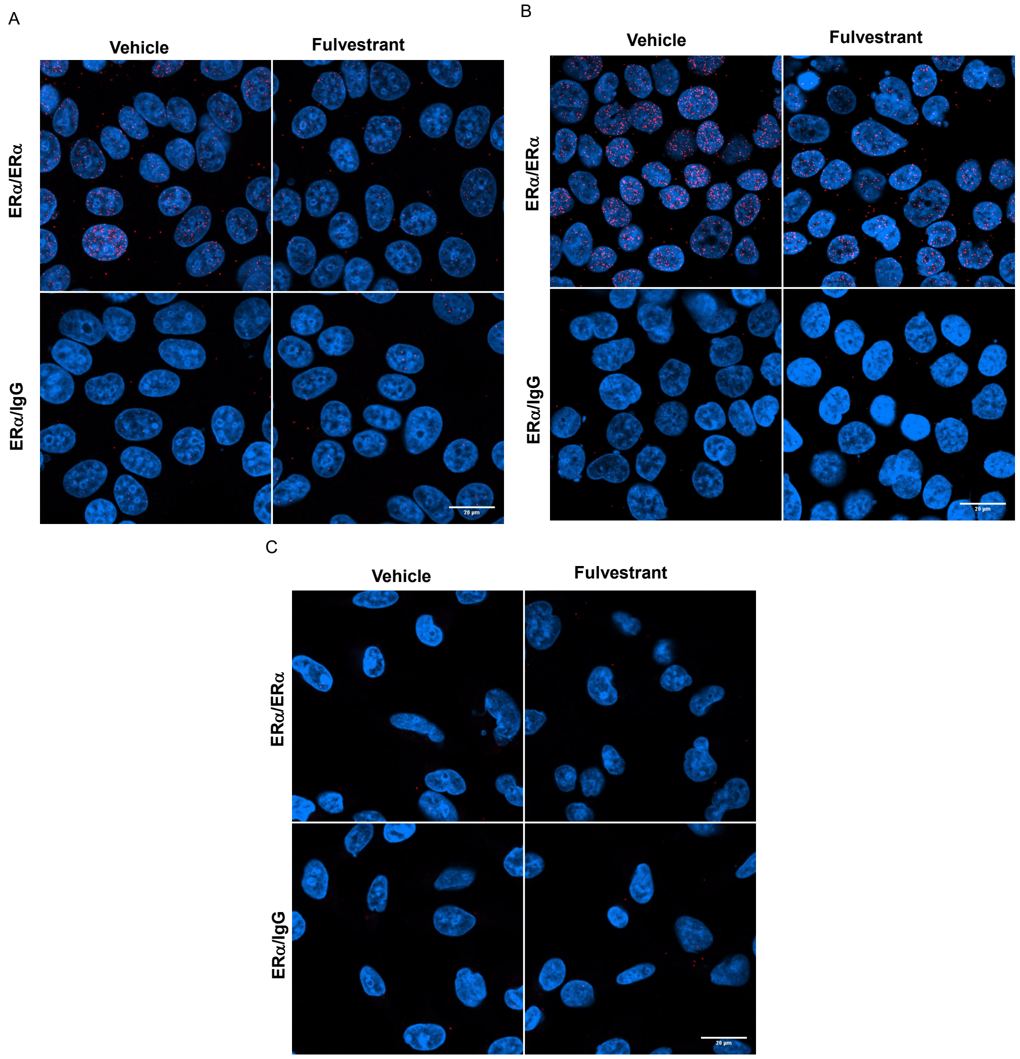


Figure S2: **Biological controls to validate ER-associated interactions in the Proximity Ligation Assay**. (A) MCF7, (B) T47D and (C) MDA-MB-231 cells were treated for 24 hours with vehicle (ethanol) or with 100nM Fulvestrant and were subsequently analysed by PLA. Dual antibody recognition of ER was used as a positive control and antibodies against ER and an isotype IgG were used together as a negative control. Fulvestrant was added as an additional biological control as it is known to degrade ER. Images were taken at 630x magnification. Cell nuclei are shown in blue and PLA signals are shown as red dots.

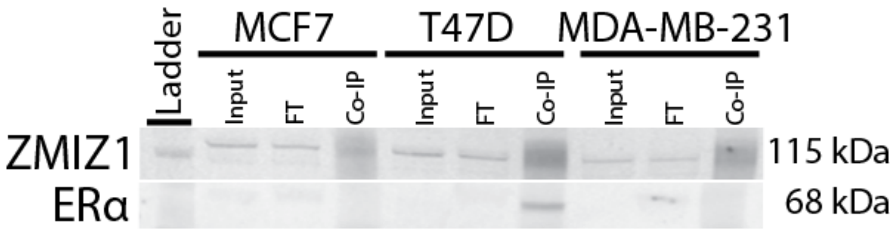


Figure S3: **Co-immunoprecipitation of Estrogen Receptor by pull-down of ZMIZ1 from cell lysate**. ZMIZ1 was detected in the Input, Flow-Through (FT) and IP for all three cell lines. The Co-IP successfully detected an interaction between ER and ZMIZ1 in the T47D cell line. The lack of detection in MCF7 can be explained by the use of native conditions without cross-linking. Cross-linking is used to enhance the detection transient interactions by qPLEX-RIME. Nonetheless, this result confirms the interaction of ZMIZ1 and ER in breast cancer cell lines models. ZMIZ1 co-IP and detection was undertaken using ab65767 (Abcam) and ER detection was undertaken with sc-8002 (Santa Cruz).

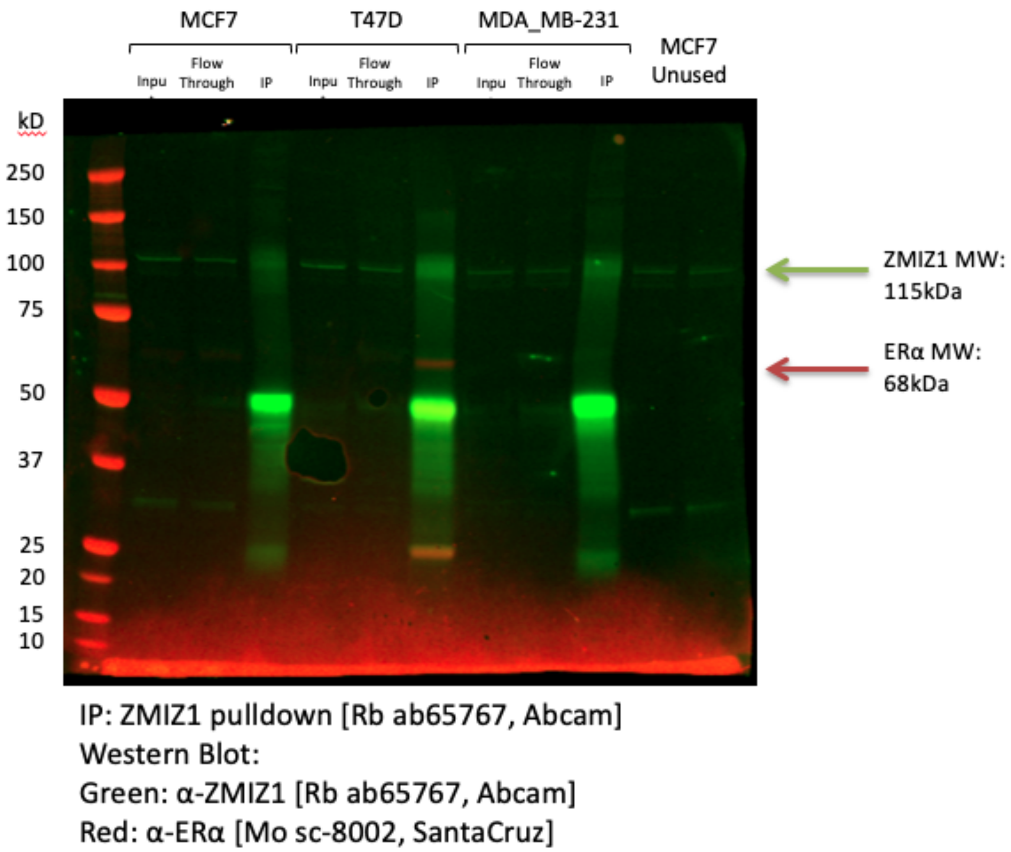


Figure S4: **Co-immunoprecipitation of Estrogen Receptor by pull-down of ZMIZ1 from cell lysate (Complete Blot)**. Nonspecific bands 50 kDa band and 17 kDa are present on the manufacture's data sheet for the ZMIZ1 antibody.

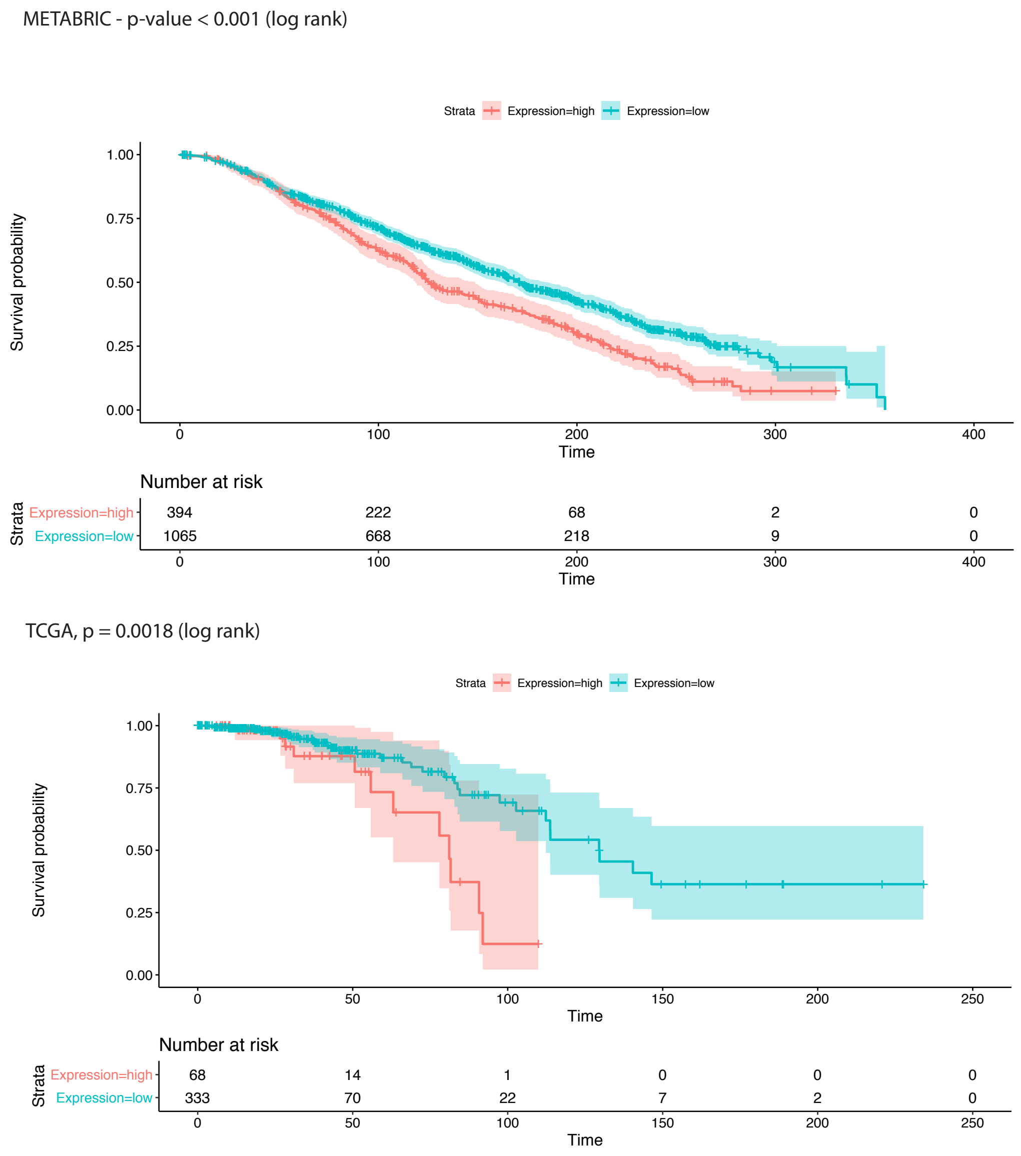


Figure S5: **ZMIZ1 is a significant predictor of survival for ER+ patients in the METABRIC and TCGA cohorts.** The METABRIC and TCGA cohorts were filtered on ER status as recorded in their respective metadata. The most suitable cut-off for expression was calculated using "survminer" package for R. In both cohorts, ZMIZ1 was a significant predictor of ER+ patient survival (METABRIC $p < 0.001$, TCGA $p = 0.0018$).

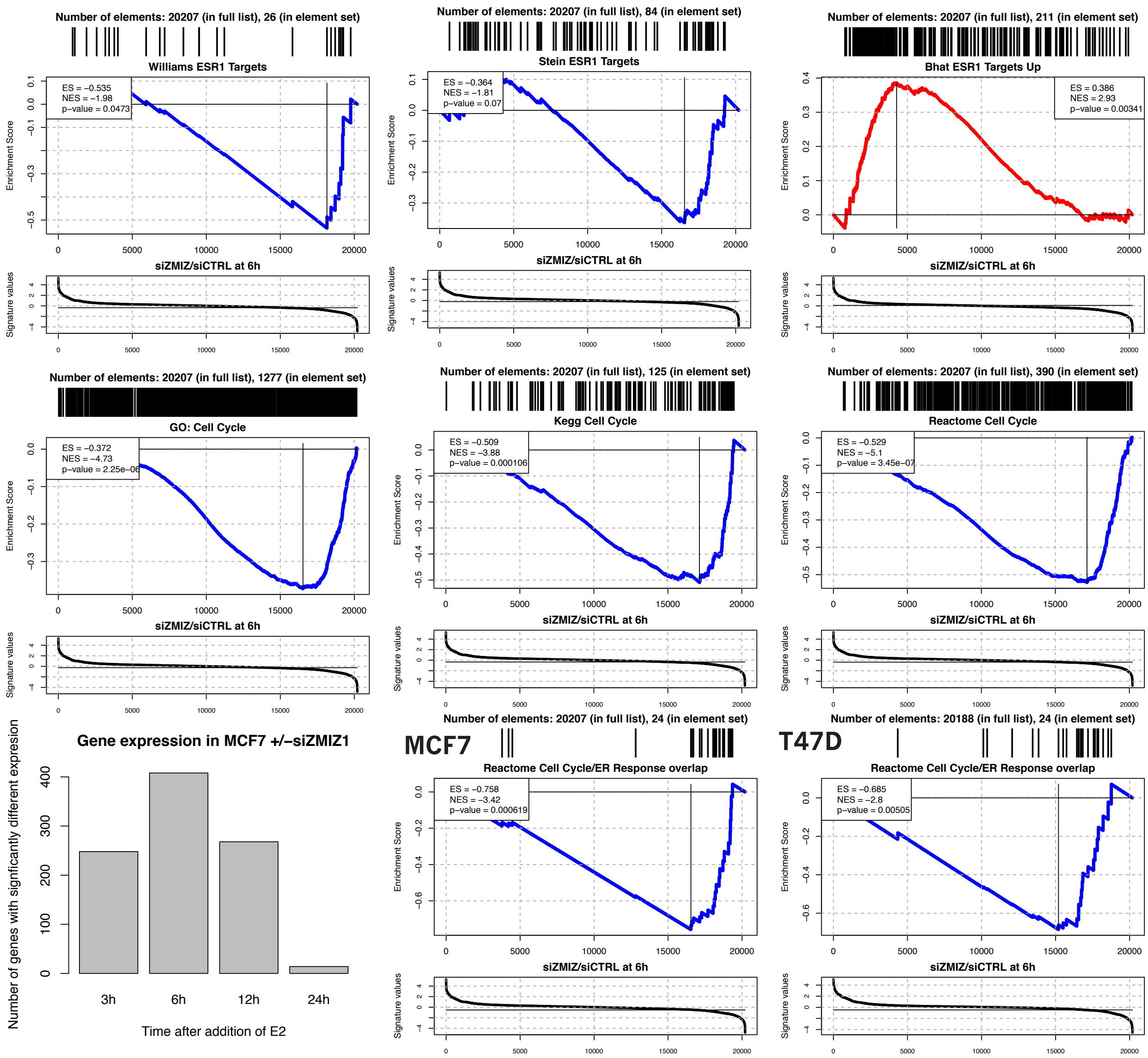


Figure S6: **ZMIZ1 knockdown delays response to E2 in ER regulated cell-cycle related genes.** GSEA analysis of RNA-seq +/-siZMIZ1. Row 1–2, GSEA of differentially expressed genes +/-siZMIZ1 at 6 hours after addition of E2. Row 3 (Left), Number of differentially expressed genes +/-siZMIZ1 at 3, 6, 12 and 24 hours after addition of E2. Row 3 (Middle and Right), GSEA of cell-cycle specific ER responsive genes shows knockdown of ZMIZ1 targets these genes.

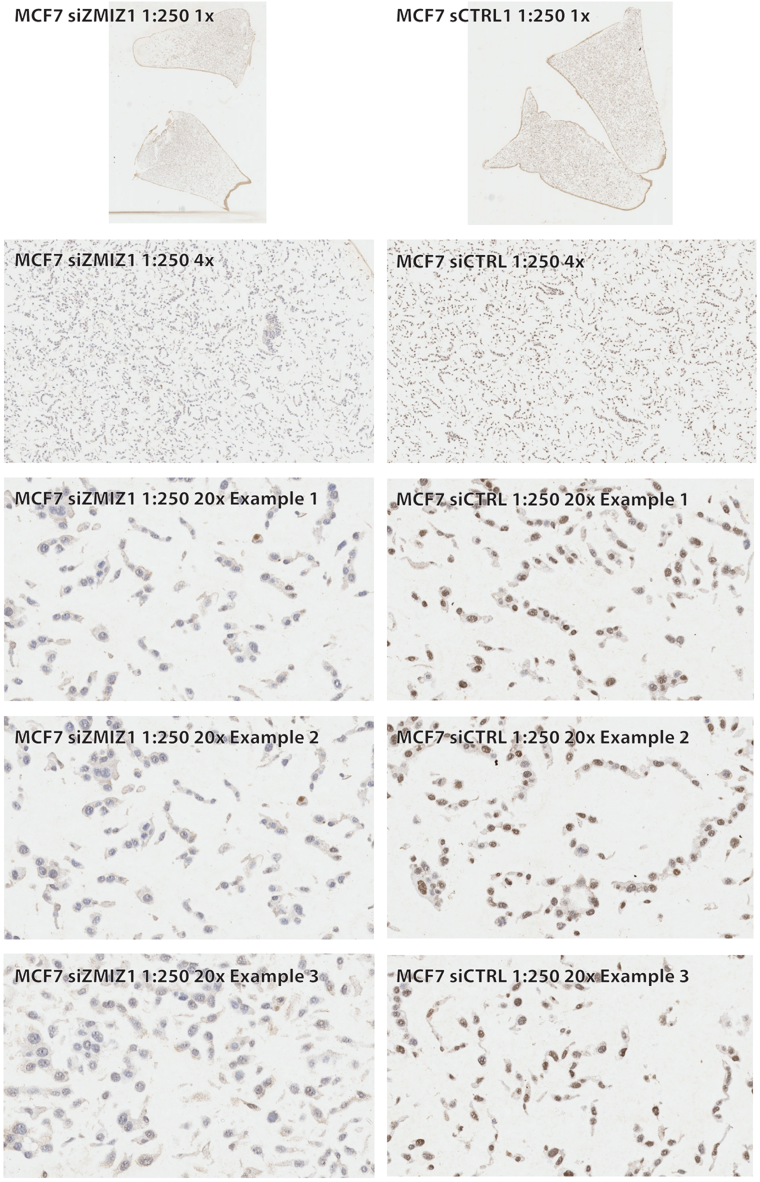


Figure S7: **ZMIZ1 Antibody validation of IHC**. Formalin fixed pellets of MCF7 cells were processed as described in the methods section. The antibody (1:250 dilution) used for the images shown was purchased from R&D Systems (AF8107) and used for all subsequent samples. Nuclear staining of ZMIZ1 was visibly reduced in the siZMIZ1 condition.
